# A one penny imputed genome from next generation reference panels

**DOI:** 10.1101/357806

**Authors:** Brian L. Browning, Ying Zhou, Sharon R. Browning

## Abstract

Genotype imputation is commonly performed in genome-wide association studies because it greatly increases the number of markers that can be tested for association with a trait. In general, one should perform genotype imputation using the largest reference panel that is available because the number of accurately imputed variants increases with reference panel size. However, one impediment to using larger reference panels is the increased computational cost of imputation. We present a new genotype imputation method, Beagle 5.0, which greatly reduces the computational cost of imputation from large reference panels. We compare Beagle 5.0 with Beagle 4.1, Impute4, Minimac3, and Minimac4 using 1000 Genomes Project data, Haplotype Reference Consortium data, and simulated data for 10k, 100k, 1M, and 10M reference samples. All methods produce nearly identical accuracy, but Beagle 5.0 has the lowest computation time and the best scaling of computation time with increasing reference panel size. For 10k, 100k, 1M, and 10M reference samples and 1000 phased target samples, Beagle 5.0’s computation time is 3× (10k), 12× (100k), 43× (1M), and 533× (10M) faster than the fastest alternative method. Cost data from the Amazon Elastic Compute Cloud show that Beagle 5.0 can perform genome-wide imputation from 10M reference samples into 1000 phased target samples at a cost of less than one US cent per sample.

Beagle 5.0 is freely available from https://faculty.washington.edu/browning/beagle/beagle.html.

## Introduction

Genotype imputation uses a reference panel of phased, sequenced individuals to estimate sequence data in target samples which have been genotyped on a SNP array.^1; 2^ Imputation of ungenotyped markers is a standard tool in genome-wide association studies because it greatly increases the number of markers that can be tested for association with a trait. Genotype imputation also provides a foundation for meta-analysis of genome-wide association studies because it converts data for samples that have been genotyped on different SNP arrays into genotype data for a shared set of sequence variants. Meta-analyses based on genotype imputation have discovered thousands of new genetic associations.^3^

The reference panel plays the primary role in determining the accuracy of imputed variants. Imputation accuracy for a variant generally increases with increasing reference panel size, and variants must be present in the reference panel in order to be accurately imputed.^4; 5^ Consequently, whenever a larger reference panel becomes available, it is advantageous to re-impute target samples with the larger panel for subsequent analysis.

The value of larger reference panels has led to steadily increasing reference panel size, with the largest reference panels to date having thousands or tens of thousands of samples.^6–12^ Ongoing, large-scale projects, such as the Trans-Omics for Precision Medicine (TopMed) program,^13^ and the National Human Genome Research Institute’s Centers for Common Disease Genomics,^14; 15^ are expected to produce reference panels with more than one hundred thousand individuals.

Increasing reference panel size increases the computational cost of imputation, and this increasing cost has motivated the development of many new computational methods and optimizations. Current imputation methods are able to make use of a rich palette of computational techniques, including the use of identity-by-descent,^6; 16^ haplotype clustering,^4; 17^ and linear interpolation^5^ to reduce the model state space, the use of pre-phasing to reduce computational complexity,^16; 18; 19^ and the use of specialized reference file formats to reduce file size and memory footprint.^4; 5; 20^ The techniques developed thus far have made it possible to provide imputation using reference panels with tens of thousands of individuals as a free web service.^4^

However, the total computational cost of imputation is substantial and increasing. Millions of individuals have been genotyped on SNP arrays and it is probable that millions more will be genotyped in the future. A sample may be re-imputed to a current state-of-the-art reference panel each time it is analyzed or included in a meta-analysis. The increasing size of reference panels increases the per-sample cost of re-imputation. If the past exponential growth in reference panel size continues, reference panels could have ten million individuals within a decade.^6; 7; 10–12^

In this paper, we present new methodology that reduces the computational cost of genotype imputation 500-fold when imputing from 10M reference samples, and we use the Amazon Elastic Compute Cloud, to show that the new methodology can impute genotypes from 10M reference samples into 1000 phased target samples for a cost of less than one US penny per sample.

The computational performance of our method is made possible by combining previous advances with several methodological improvements and innovations. Our imputation method uses a Li and Stephens haplotype frequency model,^2; 21^ with a highly parsimonious model state space that is a fraction of the size of the state space of the full LI and Stephens model. We employ a pre-processing step to reduce the full reference panel into a small number of composite reference haplotypes, each of which is a mosaic of reference haplotypes. In addition, we decouple probability calculations at genotyped and ungenotyped markers, which enables us to further reduce the number of state probabilities that must be calculated and stored.

We also make use of recent advances in simulation methods^22^ and inference of historical effective population size^23^ to generate a simulated data set that is much larger and more realistic than simulated data used in previous work.^5^ In large reference panels, many markers will be multi-allelic, and our simulation allows for multi-allelic markers, recurrent mutation, and back mutation. We simulate 10M reference samples and 1000 target samples from a population whose growth rates and demographic parameters are modeled on the UK European population.

Our new imputation method is freely available and implemented in the open source Beagle 5.0 software package.

## Methods and Materials

### Imputation Methods

We assume that the reference and target genotypes are phased and non-missing. This reduces the computational complexity of imputation from quadratic to linear in the size of the reference panel, and it simplifies the imputation problem to imputing missing alleles on a haplotype.^18^

We use the term "reference markers" to refer to markers which are genotyped in the reference panel, the term "target markers" to refer to the markers which are genotyped in the target samples, and the term "imputed markers" to refer to markers in the reference panel which are not genotyped in the target samples. We assume that the target markers are a subset of the reference markers.

Genotype imputation is based on identity by descent (IBD). Two chromosome segments that are inherited from a common ancestor without recombination since the common ancestor are said to be inherited identical by descent. In an IBD segment, the two chromosomes will have identical allele sequences except at sites which have mutated in one of the lineages since the common ancestor. We can use the genotypes at the target markers to identify long IBD segments that a target haplotype shares with the reference haplotypes. If an IBD segment is accurately identified, the ungenotyped alleles IBD segment in the target haplotype can be copied from the IBD segment in the reference haplotype. Since there is uncertainty in inferring IBD, a probabilistic model is used to account for the uncertainty and to produce a posterior probability for each possible allele at an imputed marker on the target haplotype. This probabilistic model is typically a hidden Markov model (HMM).^24^

### Hidden Markov model

Our imputation method is based on the Li and Stephens^21^ HMM. Since the Li and Stephens model has been described many times,^1; 2; 4; 5; 25^ we provide only a brief description here.

In the Li and Stephens model, the HMM state space is a matrix of reference alleles, whose rows are reference haplotypes and whose columns are markers. Each HMM state is an entry of the matrix defined by the reference haplotype (row) and marker (column), and each HMM state is labeled with the allele carried by the reference haplotype at the marker.

In imputation with the Li and Stephens model, we assume that the target haplotype corresponds to an unobserved path through the HMM state space from the first marker to the last marker that includes one state at each marker. The HMM and the observed data on the target haplotype determine a probability distribution on the paths. The HMM forward-backward algorithm^24^ is used to calculate the probability that the unobserved path passes through a HMM state (the state probability). At each marker, the sum of the probabilities of the states labeled with an allele is the imputed probability for that allele.

A HMM is defined by its state space, initial probabilities, transition probabilities and emission probabilities. The Li and Stephens HMM state space is the set of all ordered pairs (*h, m*) whose first element is a reference haplotype and whose second element is a reference marker. We label the reference haplotypes, *H*, with indices 1, 2,…, |*H*|, and we label the list of reference markers in chromosome order, *M*, with 1, 2,…, |*M*|, where |·| denotes the size of the list.

We assign an initial probability of *1/|H|* to each state at the first marker. State transitions are permitted from any state at a marker to any state at the next marker. In the Li and Stephens model used by Impute^2^ and Beagle,^5^ the state transition probabilities between marker *m −* 1 and marker *m* are defined in terms of the effective population size *N_e_*, the genetic distance *d_m_* in Morgans between markers *m −* 1 and *m*, and the number of reference haplotypes |*H*|. If the HMM state at marker *m −* 1 is on reference haplotype *h* then with probability 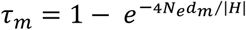, a historical recombination event will cause the HMM state at marker *m* to be randomly chosen from the *H* reference haplotypes.^2^ Otherwise (with probability 1 − *τ_m_*) the HMM state at marker *m* is required to stay on the same reference haplotype. Consequently, the probability that a state (*h, m −* 1) transitions to state (*h, m*) is (1 − *τ_m_*) *+ τ_m_/|H|*, and the probability that the state (*h,m* − 1) transitions to state (*h’, m*) with *h’ ≠ h* is *τ_m_/|H|*.

We define emission probabilities in terms of an error rate *ε* (0.0001 by default). Each state (*h, m*) emits the allele carried by reference haplotype *h* at marker *m* with probability (1 − *ε*), and emits a different allele with probability *ε*.

Our method uses several modifications of this basic Li and Stephens HMM that are used in the Beagle 4.1 imputation method: aggregate markers, restriction of the HMM to the target markers, and a sliding markers window.^5^ We briefly describe these modifications next.

As in Beagle 4.1, we collapse tightly-linked target markers within 0.005 cM into a single aggregate marker whose alleles are the sequences of alleles at the constituent markers. For an aggregate marker comprised of *l* markers, the genetic map position is the mean genetic position of the first and last markers, and the probability that a HMM state emits a different haplotype is *lε* since the emission of a different allele at any of the *l* constituent markers will cause a different haplotype to be emitted.^5^

We initially include only target markers in the HMM. After computing state probabilities at the target markers, we use linear interpolation on genetic distance to estimate HMM state probabilities at each imputed marker.^5^

Finally, we use a long sliding marker window (40 cM by default), with substantial overlap between adjacent windows (4 cM by default). Markers in the overlap between two adjacent windows will be imputed twice (once in each window). Since imputation accuracy is expected to increase with distance from a window boundary, we discard the imputed value from the window in which the position is closer to the window boundary.

### Computational Methods

Beagle 5.0 also incorporates three additional computational innovations beyond those inherited from Beagle 4.1: composite reference haplotypes, imputation on output, and an improved reference file format (binary reference format version 3).

#### Composite reference haplotypes

One approach to reducing computation time for imputation is to reduce the size of the HMM state space by using only a subset of reference haplotypes when imputing a target haplotype.^16; 19^ If the subset contains the reference haplotypes that share the longest IBD segments with the target haplotype at each position, then the subset can yield the same imputation accuracy as the full set of reference haplotypes.^6^

Previous methods for selecting a target-specific set of reference haplotypes^6; 16^ have required pre-specifying a short marker window (typically with length < 5 cM). If the marker window is too long, computational efficiency is reduced because the size of the subset must be increased in order to contain the additional long IBD segments in the larger window. However if the marker window is too short, imputation accuracy is reduced because IBD segments are truncated by the window boundaries. We present an example of this loss of accuracy in the Results section.

We have developed a method which allows imputation to use long marker windows and a small HMM state space. Instead of a using a target-specific set of reference haplotypes to construct the HMM state space, we use a target-specific set of composite reference haplotypes, where each composite reference haplotype is a mosaic of reference haplotype segments (Figure 1).

**Figure 1.**
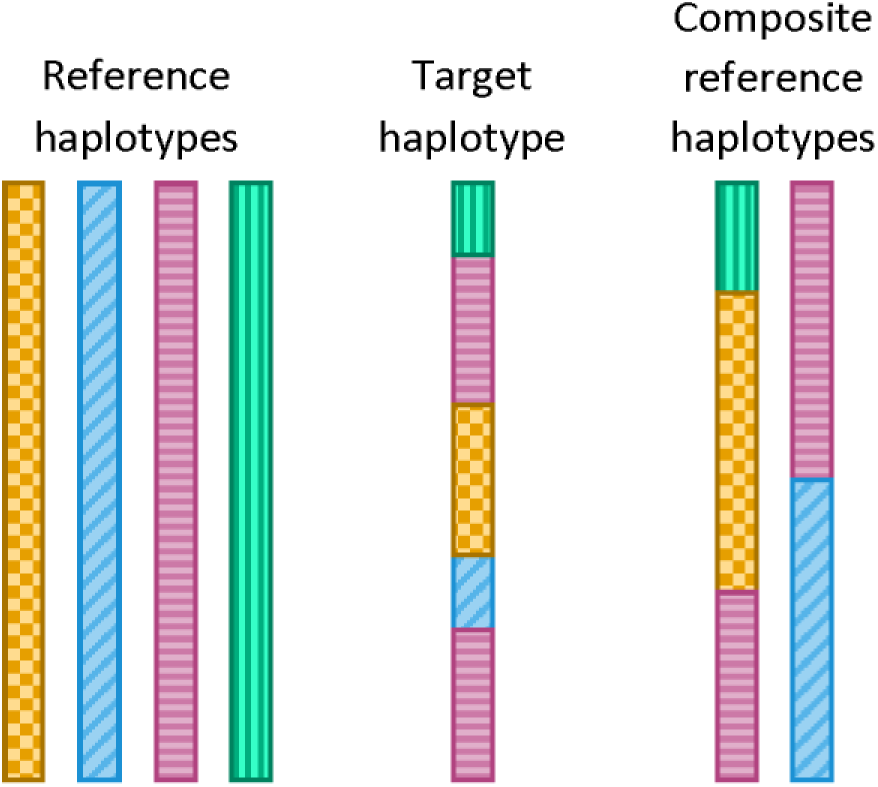
Composite reference haplotypes. Long haplotype segments with identical allele sequences are shown with the same color and pattern. The target haplotype shares five segments of identical alleles with the reference haplotypes. The two composite reference haplotypes are each a mosaic of reference haplotypes, with the mosaics chosen so that the target haplotype also shares five segments of identical alleles with the composite reference haplotypes. This permits the two composite reference haplotypes to be used in place of the four original reference haplotypes.

By construction, each haplotype segment in a composite reference haplotype will contain a long interval in which the segment and the target haplotype have identical alleles at the target markers. These intervals are called identity by state (IBS) segments. One can generally expect each long IBS segment to contain at least one long IBD segment. A relatively small number of composite reference haplotypes can be used with large marker windows because each composite reference haplotype contains many long IBS segments.

We give an informal description of composite reference haplotype construction followed by a more precise description using pseudocode. There are a fixed number of composite reference haplotypes (1600 by default), and we construct these composite reference haplotypes one haplotype segment at a time as we work through the marker window in chromosome order. As each segment is identified, it is added to one of the composite reference haplotypes. As we work through the marker window we keep track of the staleness of each composite reference haplotype, which is the distance between the current position and the composite reference haplotype’s last IBS segment. When we add a new haplotype segment, we add it to the most stale composite reference haplotype.

#### Constructing composite reference haplotypes

In order to ensure that the haplotype segments in the composite reference haplotypes cover the entire marker window and to facilitate parallel computation, we divide the marker window into consecutive, non-overlapping intervals *I*_1_*,I*_2_*,…,I_K_* of fixed length (0.1 cM length by default). In each interval *I_k_* we select a set *S_k_* of reference haplotypes that are identical by state with the target haplotype. Each *S_k_* can contain at most *s* elements (we describe how *s* is determined in the Appendix). We want to preferentially select haplotypes for *S_k_* that have longer IBS segments, and we do this by considering seven (by default) consecutive intervals beginning with *I_k_*. We add reference haplotypes to *S_k_* that are IBS with the target haplotype in seven consecutive intervals, six consecutive intervals, and so on, until *S_k_* has size *s* or until we have added all haplotypes that are identical by state in *I_k_*. All the reference haplotypes and associated IBS segments in the *S_k_* will be included in the composite reference haplotypes.

Pseudocode for construction of the composite reference haplotypes is given in Figure 2. In the pseudocode, the segments comprising each composite reference haplotype are defined by a list of starting markers *m_i_* and a list of haplotypes *h_i_*. The *i*-th segment will copy haplotype *h_i_*, starting with marker *m_i_*, and ending with marker *m_i+_*_1_ − 1 (or ending with the last marker in the window if the *i*-th segment is the last segment). We add one segment at a time by adding a starting marker *m* and a haplotype *h* to the starting marker and haplotype lists of a composite reference haplotype.

**Figure 2.**
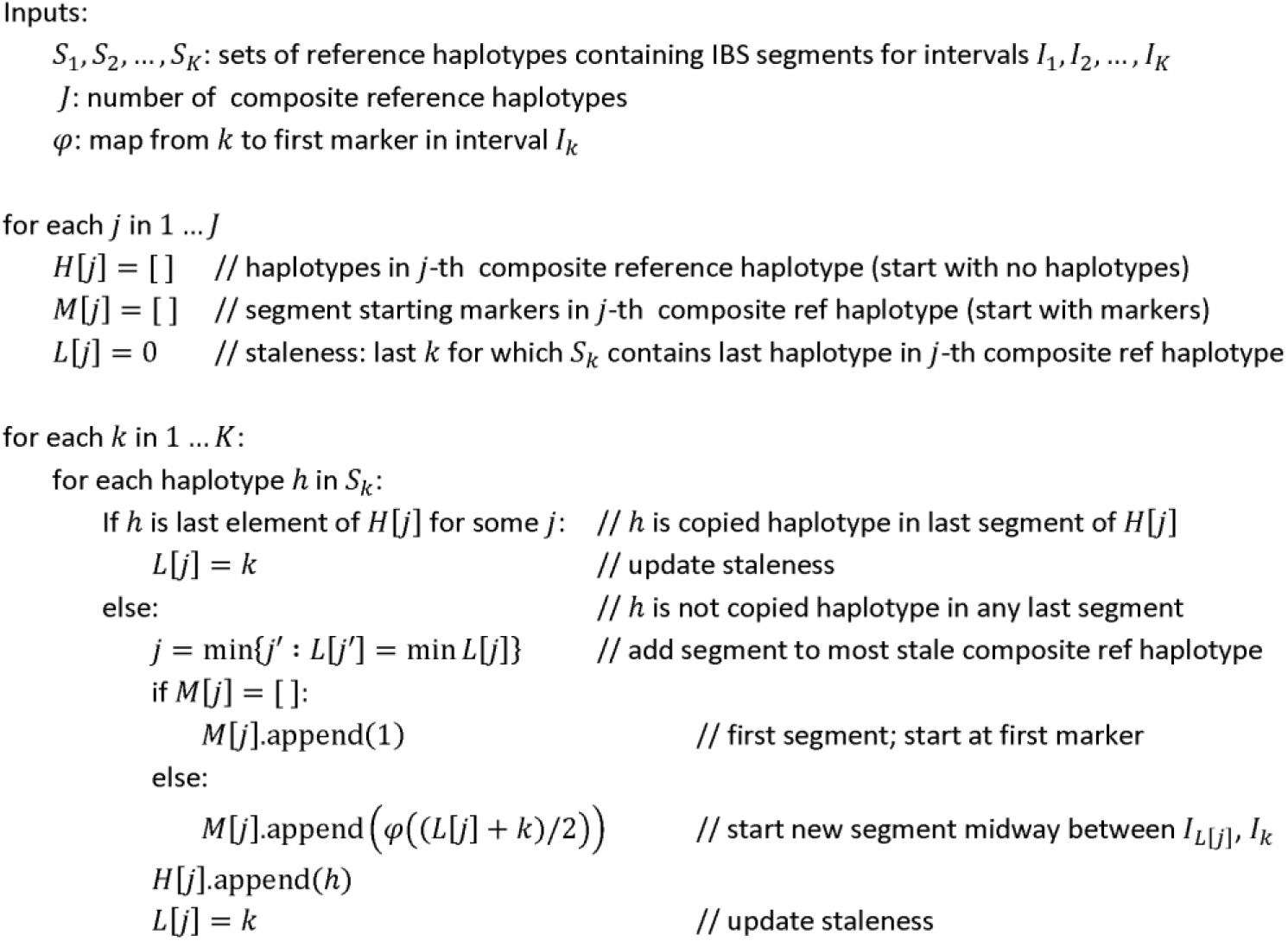
Pseudocode for constructing composite reference haplotypes

In the pseudocode, *L*[*j*] keeps track of the staleness of the *j*-th composite reference haplotype. *L*[*j*] stores the last *S_k_* which contained the most recent haplotype added to the *j*-th composite reference haplotype. If *I_k_* is the current interval being processed, and we want to add a new segment to the *j*-th composite reference haplotype, we set the starting point of the segment to a position approximately mid-way between intervals *I_L_*_[_*_j_*_]_ and *I_k_*.

#### Imputation on output

Our previous genotype imputation method (Beagle 4.1) calculates imputed allele probabilities for each individual immediately after calculating HMM state probabilities at the target markers.^5^ Beagle 5.0 postpones calculation of imputed allele probabilities until the output VCF file is constructed. This reduces memory requirements because imputation can be performed in short chromosome intervals with the results immediately written to the output VCF file. In our experience, the reduced memory footprint has the added benefit of reducing computation time.

As in Beagle 4.1, we estimate the HMM state probabilities at imputed markers using the two HMM state probabilities at the bounding target markers and linear interpolation on genetic distance, and we combine all reference haplotypes that have identical allele sequences in the interval before performing linear interpolation.^5^ Once the state probabilities are interpolated, the posterior probability for a particular allele is the sum of the state probabilities at the marker for the reference haplotypes that carry the allele. The key point is that the state probabilities at all imputed markers between two genotyped markers are obtained from the state probabilities at the bounding genotyped markers. Consequently, by storing state probabilities at consecutive markers, we can perform imputation as the VCF output file is constructed.

In practice, only a small subset of the state probabilities at consecutive markers needs to be stored. We store state probabilities at consecutive markers for a reference haplotype only if one of the state probabilities is > 1/*J*, where *J* is the number of composite reference haplotypes, or equivalently, the number of HMM states at the marker. If state probabilities at consecutive markers are not stored for a reference haplotype, they are assumed to be 0 when performing imputation.

#### Binary reference format version 3

Previously we developed a specialized file format for reference panel data called binary reference (bref) format.^5^ In bref format, a chromosome is broken into consecutive, non-overlapping intervals. If the non-major alleles of a marker are rare, bref format stores the indices of the haplotypes that carry each non-major allele. For all remaining markers in an interval, bref stores the distinct allele sequences and a pointer from each haplotype to the allele sequence carried by the haplotype at these markers. This reduces memory requirements because the number of distinct allele sequences is typically much less than the number of haplotypes. It also facilitates identification of reference haplotypes that have identical allele sequences because any two haplotypes that have a pointer to the same allele sequence will have identical alleles at all markers in the allele sequence.

Beagle 5.0 uses a new version of bref format (bref3) that reduces imputation time relative to the preceding version (bref v2). Bref3 uses two bytes to store a haplotype’s allele sequence index in a chromosome interval (bref v2 uses one byte). This permits bref3 to store allele sequences that extend over much longer chromosome intervals, which provides greater compression for large reference panels. In addition, bref3 does not gzip-compress the binary reference file (bref v2 files are gzip-compressed). This eliminates the computation time required for gzip decompression.

Table 1 reports file size for four reference file formats: gzip-compressed vcf, gzip-compressed m3vcf,^4^ bref v2,^5^ and bref3 for 10k, 100k, 1M, and 10M simulated reference UK European reference samples. These results show that bref3 scales better than bref v2 and gzip-compressed m3vcf with increasing reference panel size, and that bref3 achieves much greater compression than gzip compression.

**Table 1.** Memory (GB) required by reference file formats. Number of gigabytes required to store 10 Mb of reference sample data for 10k, 100K, 1M, and 10M simulated UK European reference samples stored in vcf.gz, m3vcf.gz, bref v2, and bref3 formats. A dash (---) indicates the compression could not be performed due to time and memory constraints.

| n | vcf.gz | m3vcf.gz | bref v2 | bref3 |
| --- | --- | --- | --- | --- |
| 10k | 0.045 | 0.011 | 0.006 | 0.017 |
| 100k | 0.51 | 0.12 | 0.064 | 0.11 |
| 1M | 7.9 | 1.9 | 0.7 | 0.7 |
| 10M | 140.0 | --- | 7.8 | 4.8 |

### Data

We compared methods using 1000 Genomes Project^11^ chromosome 20 data, Haplotype Reference Consortium^12^ chromosome 20 data, and simulated data for 10k, 100k, 1M, and 10M reference samples. Summary statistics for the six reference panels are given in Table 2.

**Table 2:**
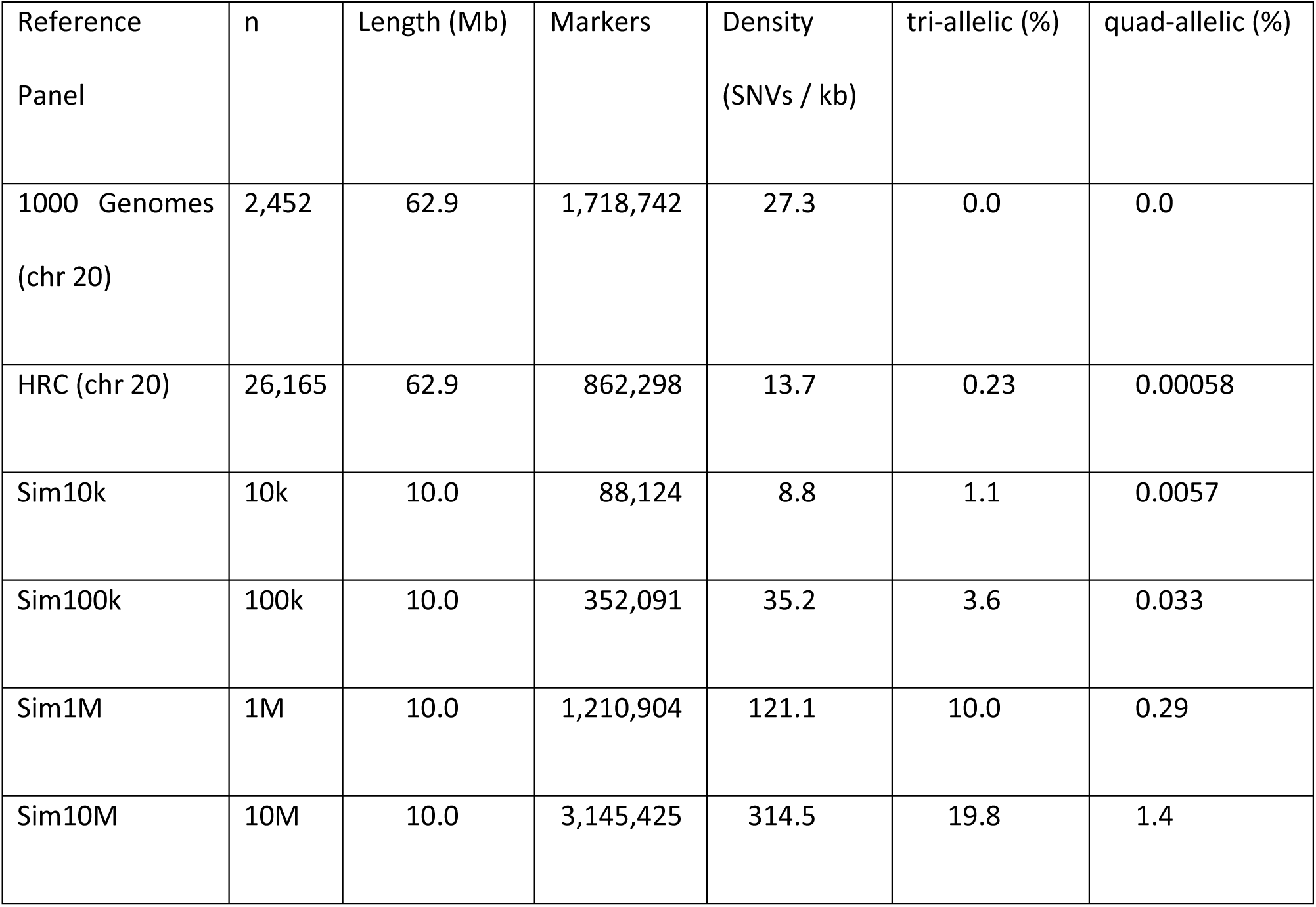
Summary statistics for reference panels used in this study.

**1000 Genomes Project data**. We downloaded the publicly available 1000 Genomes Project phase 3 version 5a data. The 1000 Genomes data contains 2504 individuals with phased sequence data from 26 populations.^11^ We randomly selected two individuals from each population (52 total) to be the imputation target. The remaining 2452 individuals were the reference panel. We restricted the 1000 Genomes reference and target data to diallelic SNVs having at least one copy of the minor allele in the reference panel. After this marker filtering there were 1,718,742 markers on chromosome 20. In the target samples, we masked chromosome 20 markers which were not on the Illumina Omni2.5 array, resulting in 54,885 target markers.

**Haplotype Reference Consortium data**. The publicly-available Haplotype Reference Consortium (HRC) data was downloaded from the European Genome-Phenome Archive^26^ at the European Bioinformatics Institute (Accession EGAD00001002729).The HRC data include 27,165 individuals with phased SNV genotypes, primarily from low-coverage sequencing. The chromosome 20 HRC data consist of 884,983 SNVs at 882,742 distinct positions. Some positions have multiple SNVs because in the publicly-available VCF file, multi-allelic markers are represented as multiple diallelic SNV markers at the same position. Since the HRC panel is predominantly European, we randomly selected 1000 individuals from the constituent UK10K study^27^ to be the imputation target. The remaining 26,165 individuals were the reference panel. We report computation time for imputation into 10, 100, and 1000 target samples. After excluding monomorphic markers in the reference samples there were 864,376 SNVs at 862,298 unique positions. In the target samples, we masked chromosome 20 markers which were not on the Omni2.5 array or which had more than one ALT allele in the reference samples, resulting in 55,013 target markers.

**Simulated data**. We used the msprime program^22^ (Supplemental Text S14) to simulate 10Mb of sequence data for 10,001,000 diploid samples with a demographic history modeled on the UK European population. Recent growth rates for the UK European effective population size were obtained from the IBDNe program.^5^ We created four reference panels with 10k, 100k, 1M, and 10M samples by selecting the first 10k, 100k, 1M, and 10M simulated samples. The last 1000 simulated samples became the imputation target. We report computation time for imputation into 10, 100, and 1000 target samples.

Since msprime simulates diallellic variants, we devised a strategy for simulating multi-allelic single nucleotide variants which permits recurrent and back mutation. When running msprime, we increased the length of the 10 Mb simulated region by a factor of 100 and decreased the mutation rate and recombination rate by a factor of 100. After generating data with msprime, we divided the simulated region into non-overlapping 100 base pair windows and collapsed the VCF records in the *k*-th window into a single genomic position whose position was *k*. In each 100 base pair window, we reconstructed the phylogenetic binary tree for the haplotypes in each window from the alleles carried by the haplotypes in the window. At each mutation event in the phylogenetic tree, we randomly chose a new allele from the 3 possible new alleles, and the haplotypes in each leaf of the tree were assigned the allele inherited by that leaf.

We randomly selected 3,333 diallelic variants having minor allele frequency ≥ 0.05 to be the genotyped markers in the imputation target samples. The resulting target marker density (1 marker per 3.3 kb) corresponds to a 1M SNP array in humans.

We also created one chromosome-scale reference panel by making 30 copies of the 10M sample reference panel, and shifting each copy by a multiple of 10 Mb. We used this large reference panel to explore the computational performance of our imputation method when imputing a long 300 Mb chromosome from 10M reference samples.

### Comparison of methods

We compared the imputation accuracy and the computation time of our imputation method with four recent versions of three published imputation methods: Beagle 4.1 (27Jan18.7e1),^5^ Impute4 r265.2,^16; 18; 19; 28^ Minimac3 v2.0.1,^4; 18^ and Minimac4 v1.0.0.^29^

We used default parameters for each program, except when otherwise noted. When running Impute4, we used the no_maf_align option because strand alignment of reference and target variants was not needed. Impute4 requires specification of the analysis region, which was the entire chromosome or 10 Mb simulated region, except for the HRC reference panel. For the HRC Impute4 analysis, we split chromosome 20 into two analysis regions with 250 kb overlap (the default Impute4 buffer parameter) in order to perform imputation within the available computer memory. For Beagle 4.1, we increased the window parameter so that the imputation analysis used a single marker window, except for the simulated 10M reference panel. For the 10M reference panel, Beagle 4.1 used a marker window of 1,400,000 markers (≈ 4.45 cM) with an overlap of 315,000 markers (≈ 1 cM) between adjacent windows in order to run within the available computer memory.

Beagle and Impute require a user-specified genetic map. We used the HapMap2 genetic map^8; 30^ for analyses with real data, and we used the true genetic map for analyses with simulated data. Minimac does not require a genetic map because recombination parameters are estimated and stored when producing the m3vcf format input file for the reference data.

Beagle 4.1, Beagle, 5.0, Minimac3, and Minimac4 were run using their specialized formats for reference panel data: bref v2^5^ for Beagle 4.1, bref3 for Beagle 5.0, and m3vcf for Minimac3 and Minimac4. We used Minimac3 to construct the m3vcf files. We succeeded in creating of an m3vcf reference file for 1M reference samples by borrowing a compute node with 1 TB of memory, but we were unable to create an m3vcf reference file for the largest 10M reference panel due to time and memory constraints.

We report the squared correlation (*r^2^*) between the true number of non-major alleles on a haplotype (0 or 1) and the posterior imputed allele probability. Since there is little information to estimate squared correlation for an individual marker when minor allele counts are low, we binned imputed minor alleles according to the minor allele count in the reference panel, and we calculated *r^2^* for the imputed minor alleles in each minor allele count bin.

All imputation analyses were run on a dedicated 12-core 2.6 GHz computer with Intel Xeon E5-2630v2 processors and 128 GB of memory, except where otherwise noted. We evaluated one program at a time using 1 and 12 computational threads. For the single-threaded analyses, we report the sum of the system and user time returned by the Unix time command. For multi-threaded analyses, we report the real (wall-clock) time returned by the Unix time command.

### Multi-allelic markers

Beagle is the only program evaluated that permits reference markers to have more than two alleles. For the other programs (Minimac and Impute), triallelic and quadallelic markers in the simulated reference panels and the 1000 Genomes reference panel were partitioned into two and three diallelic markers respectively using the bcftools norm -m command (see electronic resources). The downloaded HRC reference panel represents multi-allelic markers as multiple diallelic markers, and was left as is.

## Results

Beagle was evaluated using all reference panels. Minimac3 and Minimac4 were run with all reference panels except the largest reference panel (10M) because an m3vcf format file could not be constructed for this reference panel. Impute4 was run with all reference panels except the three largest reference panels (100k, 1M, and 10M). The Impute4 error message for the 100k reference panel indicated that Impute4 is limited to 65,536 reference haplotypes.

### Accuracy

All the genotype imputation methods that we evaluated are based on the Li and Stephens probabilistic model^31^ and have essentially the same imputation accuracy (Figure 3), which is consistent with previous reports.^4; 5^ The apparent difference in accuracy between Beagle 4.1 and Beagle 5.0 with the 10M reference panel is due to Beagle 4.1 using a short (≈4.45 cM) marker window in order to run within the available computer memory. Experiments with different window and overlap lengths, showed that the difference in imputation accuracy for the 10M reference panel is almost entirely explained by the difference in window length (Figure S13).

**Figure 3.**
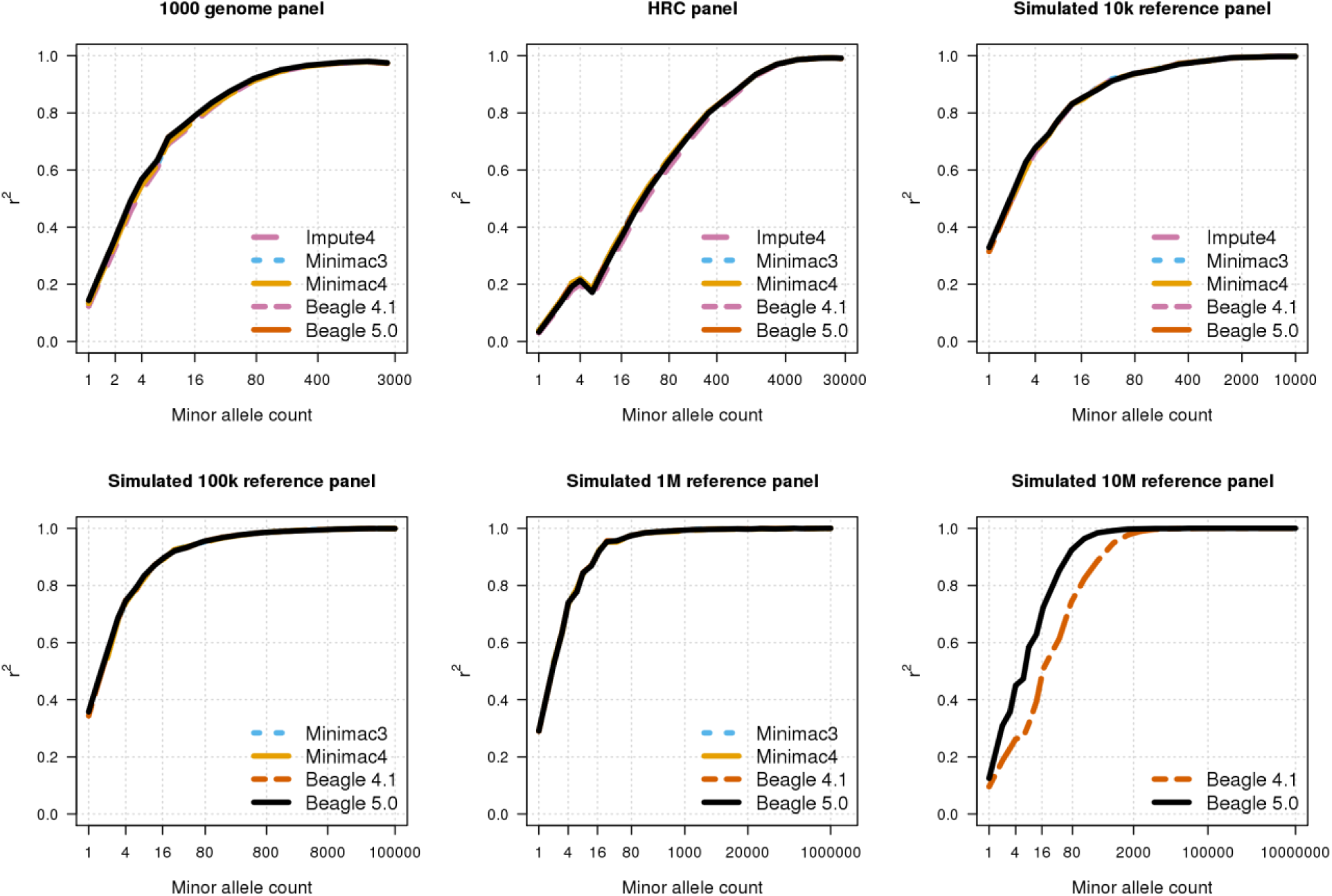
Genotype imputation accuracy. Genotype imputation accuracy when imputing genotypes from a 1000 Genomes Project reference panel (n=2,452), a Haplotype Reference Consortium reference panel (n=26,165), and from 10k, 100k, 1M, and 10M simulated UK-European reference samples. Imputed alleles are binned according to their minor allele count in each reference panel. The squared correlation (*r^2^*) between the true number of alleles on a haplotype (0 or 1) and the imputed posterior allele probability is reported for each minor allele count bin. The horizontal axis in each panel is on a log scale. The difference in accuracy for 10M reference samples is due to a difference in length of marker window.

### Single-threaded computation time

Single-threaded computation times for 10k, 100k, 1M, and 10M reference samples and 1000 target samples are shown in Figure 4. Beagle 5.0’s computation time was 3 times (10k), 12 times (100k), 43 times (1M), and 533 times (10M) faster than the fastest alternative method. For the 1000 Genomes reference panel with 52 target samples, Beagle 5.0 was 1.6× faster than the fastest alternative method (Table S1). For the HRC reference panel with 1000 target samples, Beagle 5.0 was 8.7× faster than the fastest alternative method (Table S2). Beagle 5.0 had the lowest computation time in all single-threaded analyses and the best scaling of computation time with increasing reference panel size.

**Figure 4.**
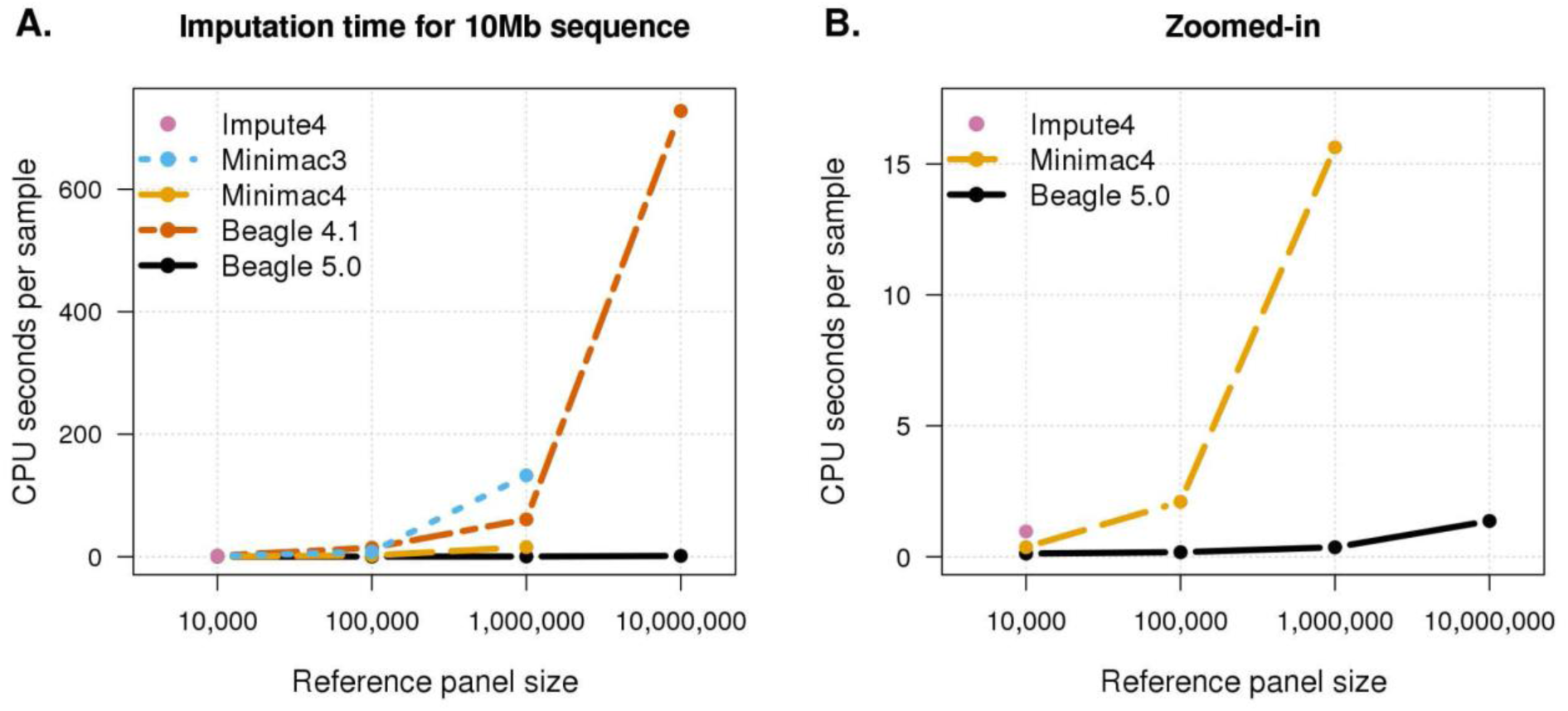
Single-threaded computation time. Per-sample CPU time when imputing a 10 Mb region from 10k, 100k, 1M, and 10M simulated UK-European reference samples into 1000 target samples using one computational thread. CPU time is the sum of the system and user time returned by the Unix time command. Impute4 was run with only the 10k reference panel due to software limitations. Minimac3 and Minimac4 were not run with the 10M reference panel due to memory and time constraints. A) Results for Impute4, Minimac3, Minimac4, Beagle 4.1 and Beagle 5.0. B) Zoomed-in results for Impute4, Minimac4, and Beagle 5.0.

Figure 4A shows that the latest versions of imputation methods have achieved substantial improvements in computational efficiency. Beagle 5.0 is significantly faster than Beagle 4.1, Minimac4 is significantly faster than Minimac3, and a comparison with imputation results in previous studies^4; 5^ for the 1000 Genomes and HRC reference panels shows that Impute4 is significantly faster than Impute2.

One striking feature of Figure 4B is the sublinear scaling of computation time with increasing reference panel size for Minimac4 and Beagle 5.0. This is particularly pronounced for Beagle 5.0. Moving from 10k to 10M reference samples increases the number of reference samples by a factor of 1000 and the number of imputed markers by a factor of 37, but Beagle 5.0’s imputation time increases by only a factor of 11 (Tables S3, S6). Sublinear scaling in reference panel size over this range of reference panel sizes is made possible by the use of specialized file formats for the reference panel (bref3 for Beagle 5.0 and m3vcf for minimac4) which scale sublinearly in reference panel size (Table 1).

### Multi-threaded computation time

Beagle and Minimac can be run with multiple threads, which reduces wall-clock computation time and permits a single copy of the reference panel data to be stored in memory and used by all CPU cores. Multi-threaded computation times for 10k, 100k, 1M, and 10M reference samples and 1000 target samples are shown in Figure 5. Beagle 5.0’s computation time was 5 times (10k), 23 times (100k), 156 times (1M), and 458 times (10M) faster than the fastest alternative method. For the 1000 Genomes reference panel, Beagle 5.0 was 3.6× faster than the fastest alternative method (Table S7). For the HRC reference panel with 1000 target samples, Beagle 5.0 was 11× faster than the fastest alternative method (Table S8). Beagle 5.0 had the lowest wall-clock time in all multi-threaded analyses and the best scaling of wall-clock time with increasing reference panel size.

**Figure 5.**
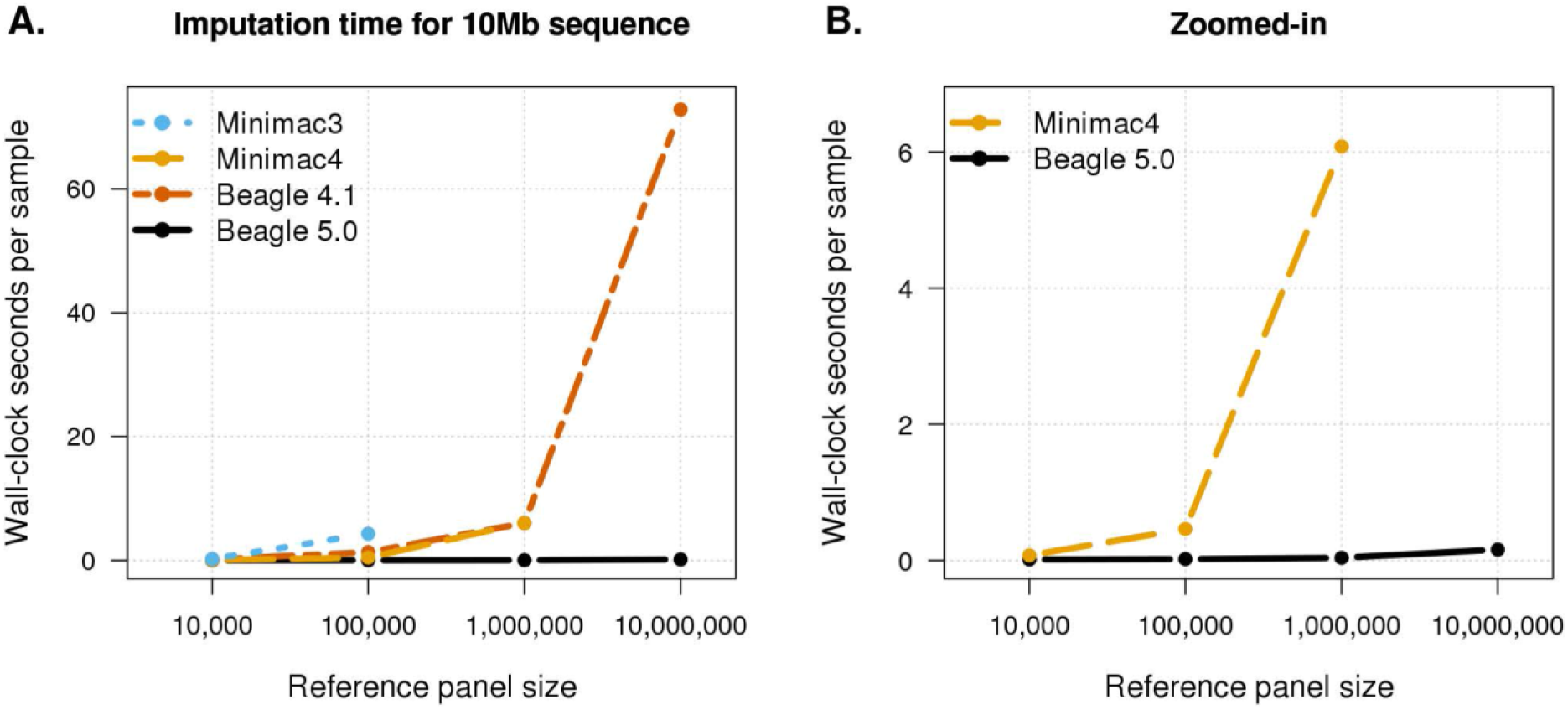
Multi-threaded computation time. Per-sample wall-clock time when imputing a 10 Mb region from 10k, 100k, 1M, and 10M simulated UK-European reference samples into 1000 target samples using 12 computational threads. Minimac3 was not run with the 1M reference panel using 12 threads due to memory constraints. Minimac3 and Minimac4 were not run with the 10M reference panel due to memory and time constraints. A) Results for Minimac3, Minimac4, Beagle 4.1 and Beagle 5.0. B) Zoomed-in results for Minimac4 and Beagle 5.0.

### Computational efficiency with decreasing target sample size

The computation time results in Figures 4 and 5 are for batches of 1000 target samples. We also examined computation times for batches of 100 and 10 target samples (Supplemental Figures S2-S6 and S8-S12). In general, the computation time per sample increases as the number of target samples decreases. This is because the time required to read reference panel data from persistent storage is independent of the number of target samples. As the target sample size decreases, this fixed computational cost is shared among fewer target samples.

As the ratio of the number of reference to target samples increases, the time required to read the reference panel into memory will eventually dominate the total computation time. For example, the computation times when imputing from 10M reference samples into 100 or 10 target samples differ by only 20% because most of the computation time in these two analyses is spent reading in the reference panel data (Table S6).

### Genome-wide imputation on the Amazon Compute Cloud

We benchmarked genome-wide imputation time and cost when imputing from 10M reference samples into 1000 target samples using the Amazon Web Services (AWS) compute cloud. We measured the cost of imputing a long 300 Mb chromosome. We used an AWS Elastic Compute Cloud (EC2) r4.4xlarge server with 16 virtual CPUs, 122 GB of memory, and a 150 GB Elastic Block Store (EBS) general purpose solid state drive. We copied data to the EBS drive via Amazon’s Simple Storage Service (S3), and we used the OpenJDK Java 1.8 runtime environment. We used a 30 cM analysis window to ensure that the memory requirements did not exceed the memory available on the EC2 server. The EC2 server ran for 97 minutes, which included 2.5 minutes for status checks and setting up the computing environment, 16 minutes for copying the program and data files onto the EBS drive, 78 minutes for genotype imputation, and 0.5 minutes for copying the imputation output files to our local computer. For comparison, we note that the same analyses can be performed with 40 cM windows on our local compute 12-core server with 128 GB of memory and a wall-clock computation time of 78 minutes.

The cost of the imputation analysis on the Amazon cloud was $0.63, which is comprised of the cost of the EC2 r4.4xlarge spot instance ($0.24), the cost of the EBS drive ($0.03), the cost of transferring data to S3 storage and briefly storing data there ($0.05), the cost to transfer 3.1 GB of imputed output data from AWS to our local computers ($0.28) and state taxes ($0.02). The per-hour cost of the EC2 r4.4xlarge spot instance was $0.15 per hour on the Ohio region servers, and this cost ranged between $0.1489 and $0.1537 per hour during a four month period from January to April, 2018. Since our analysis imputed one-tenth of the human genome, we extrapolate a total cost of $6.30 for imputation of 1000 target samples, or equivalently a genome-wide imputation cost of less than one penny per sample when imputing from 10M reference samples into 1000 target samples with Beagle 5.0.

## Discussion

We have presented a new genotype imputation method implemented in Beagle 5.0 which has similar accuracy and much faster computation time than the Beagle 4.1,^5^ Impute4,^28^ Minimac3,^4^ and Minimac4^29^ imputation methods. The Beagle 5.0 method had the lowest computation time for all reference panel sizes (2.5k - 10M) and target sample sizes (10–1000) considered. Imputation of one tenth of the genome from a reference panel with 10M individuals and one variant per 3 base pairs was performed on the Amazon Web Services compute cloud for $0.00063 per sample, which corresponds to a genome-wide imputation cost of 0.63 cents per sample.

The computational efficiency of Beagle 5.0 is due to several methodological improvements and innovations. The use of composite reference haplotypes permits modeling of arbitrarily large genomic regions with a parsimonious statistical model. An improved reference file format (bref3) reduces the computation time required to read in large reference panels relative to Beagle 4.1. Delaying imputation until output file construction reduces memory use because it permits imputation to be performed incrementally in short genomic intervals with the imputed data written immediately to the output VCF file.

As the size of the reference panel increases, the number of multi-allelic variants in the reference panel also increases. Many genotype imputation programs require multi-allelic markers to be represented as multiple diallelic markers. However, splitting a multi-allelic marker into diallelic markers is problematic because the constraint that posterior allele probabilities must sum 1.0 cannot be enforced, potentially resulting in a loss of accuracy and inconsistent imputed data. Among the imputation methods evaluated in this study, Beagle was the only method that does not require multi-allelic markers to be represented as multiple diallelic markers.

The Beagle 5.0 genotype imputation method effectively solves the problem of imputing SNV variants in large batches of target samples. However, there is scope for further methodological work to improve computational efficiency when imputing from large reference panels into small batches of target samples, and in the development and evaluation of imputation methods for non-SNV variants, such as HLA alleles,^32–35^ STR alleles,^36^ and other structural variants.

When reference panel size is much greater than the target panel size, the time required to read the reference panel from persistent storage dominates the computation time. For a fixed size reference panel, we observed a large increase in Beagle 5.0’s imputation time per sample when the reference panel size is 1000 or 10,000 times larger than the target sample size (Tables S3-S6). Like many imputation programs, Beagle 5.0 requires all target samples in the same input file to be genotyped for the same set of markers, which limits the number of target samples that can be included in an imputation analysis. One potential approach to reducing imputation time for a small number of target samples would be to modify imputation software to allow multiple input target files to be specified so that the time required to read in the reference panel data can be shared among a larger number of target samples. Although an individual researcher may not have a large number of target samples, an imputation server^4^ could use this approach to impute multiple batches of target samples submitted by different researchers in a single analysis.

The computation time for imputation with Beagle 5.0 scales sublinearly with reference panel size over the range of reference panel sizes included in this study. A 1000-fold increase in the number of reference samples and a 37-fold increase in the number of imputed markers led to only an 11-fold increase in computation time. Although we have limited our evaluation to reference panels with at most 10M individuals, these results suggest that Beagle 5.0 can analyze much larger reference panels than analyzed here.

## Appendix: The maximal size of *S_k_*

We enforce a maximal size, *s*, for each *S_k_* set in order to ensure that each segment in a composite reference haplotype exceeds a minimum length L cM. If the intervals *I_k_* used to define the *S_k_* have length *I* cM, there are *L/I* intervals in *L* cM, and the *S_k_* sets for these intervals will contain ≤ *sL/I* distinct haplotypes. If there are *J* composite reference haplotypes, we set *s* to be the largest integer such that *sL/I < J*. This ensures the number of haplotype segments added in *L* cM is less than the number of composite reference haplotypes. Consequently, each segment must be at least *L* cM in length. By default there are *J* = 1600 composite reference haplotypes, the interval length is *I* = 0.1, and the minimum segment length is *L =* 6 cM, so that *s* = [*JI/L*] *=* 26.

## Supplemental Data

Supplemental Data include twelve tables, one figure, and msprime code for simulating the genetic data.

## Acknowledgements

Research reported in this publication was supported by the National Human Genome Research Institute of the National Institutes of Health under Award Number R01HG008359. The content is solely the responsibility of the authors and does not necessarily represent the official views of the National Institutes of Health.

## Web resources

Beagle 5.0 and bref3 programs, http://faculty.washington.edu/browning/beagle/beagle.html

The 1000 Genomes Project Phase 3 version 5a data, ftp://ftp.1000genomes.ebi.ac.uk/vol1/ftp/release/20130502/

Haplotype Reference Consortium data, https://www.ebi.ac.uk/ega/datasets/EGAD00001002729

bcftools 1.5 http://www.htslib.org/doc/bcftools.html

## Supplemental Data

### Single-threaded imputation times

**Table S1:** Single-threaded CPU time for imputation from 2,452 reference samples from the 1000 Genomes Project

**Table S2:** Single-threaded CPU time for imputation from 26,165 Haplotype Reference Consortium reference samples

**Table S3:** Single-threaded CPU time for imputation from 10k simulated UK-European reference samples

**Table S4:** Single-threaded CPU time for imputation from 100k simulated UK-European reference samples

**Table S5:** Single-threaded CPU time for imputation from 1M simulated UK-European reference samples

**Table S6:** Single-threaded CPU time for imputation from 10M simulated UK-European reference samples

### Multi-threaded imputation times

**Table S7:** Multi-threaded wall-clock time for imputation from 2,452 reference samples from the 1000 Genomes Project

**Table S8:** Multi-threaded wall-clock time for imputation from 26,165 Haplotype Reference Consortium reference samples

**Table S9:** Multi-threaded wall-clock time for imputation from 10k simulated UK-European reference samples

**Table S10:** Multi-threaded wall-clock time for imputation from 100k simulated UK-European reference samples

**Table S11:** Multi-threaded wall-clock time for imputation from 1M simulated UK-European reference samples

**Table S12:** Multi-threaded wall-clock time for imputation from 10M simulated UK-European reference samples

### Effect of window length and overlap on imputation accuracy

**Figure S13:** Effect of window length and overlap on imputation accuracy

### Msprime simulation script

**Text S14:** Msprime script for simulating UK-European samples

**Table S1:** Single-threaded CPU time for imputation from 2,452 reference samples from the 1000 Genomes Project. Single-threaded CPU time for Beagle 5.0, Beagle 4.1, Minimac4, Minimac3, and Impute4 to impute chromosome 20 (1,718,742 markers) from 2,452 reference samples from the 1000 Genomes Project into 52 target samples genotyped for 55,885 markers. Imputation analyses were run on a 2.6 GHz Intel Xeon E5-2630v2 computer with 128 GB of memory. CPU time is the sum of user and system time reported by the Unix time command. Relative CPU time is the ratio of the CPU time to the Beagle 5.0 CPU time.

| Method | Target (n) | CPU seconds per target sample | Relative CPU time |
| --- | --- | --- | --- |
| Beagle 5.0 | 52 | 2.57 | 1.00 |
| Beagle 4.1 | 52 | 6.58 | 2.56 |
| Minimac4 | 52 | 3.97 | 1.55 |
| Minimac3 | 52 | 8.57 | 3.34 |
| Impute4 | 52 | 7.99 | 3.11 |

**Table S2:** Single-threaded CPU time for imputation from 26,165 Haplotype Reference Consortium reference samples. Single-threaded CPU time for Beagle 5.0, Beagle 4.1, Minimac4, Minimac3, and Impute4 to impute chromosome 20 (862,298 markers) from 26,165 Haplotype Reference Consortium reference samples into 1000, 100, and 10 target samples genotyped for 55,013 markers. Imputation analyses were run on a 2.6 GHz Intel Xeon E5-2630v2 computer with 128 GB of memory. CPU time is the sum of user and system time reported by the Unix time command. Relative CPU time is the ratio of the CPU time to the Beagle 5.0 CPU time for the same number of target samples.

| Method | Target (n) | CPU seconds per target sample | Relative CPU time |
| --- | --- | --- | --- |
| **Beagle 5.0** | 1000 | 1.79 | 1.00 |
| **Beagle 4.1** | 1000 | 40.99 | 22.95 |
| **Minimac4** | 1000 | 15.91 | 8.91 |
| **Minimac3** | 1000 | 15.59 | 8.73 |
| **Impute4** | 1000 | 39.84 | 22.31 |
| **Beagle 5.0** | 100 | 2.34 | 1.00 |
| **Beagle 4.1** | 100 | 44.82 | 19.13 |
| **Minimac4** | 100 | 20.44 | 8.72 |
| **Minimac3** | 100 | 16.01 | 6.83 |
| **Impute4** | 100 | 44.05 | 18.80 |
| **Beagle 5.0** | 10 | 7.24 | 1.00 |
| **Beagle 4.1** | 10 | 88.67 | 12.25 |
| **Minimac4** | 10 | 66.98 | 9.25 |
| **Minimac3** | 10 | 19.85 | 2.74 |
| **Impute4** | 10 | 87.75 | 12.12 |

**Table S3:** Single-threaded CPU time for imputation from 10k simulated UK-European reference samples. Single-threaded CPU time for Beagle 5.0, Beagle 4.1, Minimac4, Minimac3, and Impute4 to impute a 10 Mb region (88,124 markers) from 10k simulated UK-European reference samples into 1000, 100, and 10 target samples genotyped for 3,333 multi-allelic markers. Imputation analyses were run on a 2.6 GHz Intel Xeon E5-2630v2 computer with 128 GB of memory. CPU time is the sum of user and system time reported by the Unix time command. Relative CPU time is the ratio of the CPU time to the Beagle 5.0 CPU time for the same number of target samples.

| Method | Target (n) | CPU seconds per target sample | Relative CPU time |
| --- | --- | --- | --- |
| Beagle 5.0 | 1000 | 0.12 | 1.00 |
| Beagle 4.1 | 1000 | 1.81 | 14.86 |
| Minimac4 | 1000 | 0.36 | 2.99 |
| Minimac3 | 1000 | 0.82 | 6.78 |
| Impute4 | 1000 | 0.96 | 7.93 |
| Beagle 5.0 | 100 | 0.20 | 1.00 |
| Beagle 4.1 | 100 | 1.99 | 10.08 |
| Minimac4 | 100 | 0.47 | 2.38 |
| Minimac3 | 100 | 0.85 | 4.32 |
| Impute4 | 100 | 1.11 | 5.62 |
| Beagle 5.0 | 10 | 0.76 | 1.00 |
| Beagle 4.1 | 10 | 3.38 | 4.42 |
| Minimac4 | 10 | 1.49 | 1.95 |
| Minimac3 | 10 | 1.15 | 1.51 |
| Impute4 | 10 | 2.55 | 3.34 |

**Table S4:** Single-threaded CPU time for imputation from 100k simulated UK-European reference samples. Single-threaded CPU time for Beagle 5.0, Beagle 4.1, Minimac4, Minimac3, and Impute4 to impute a 10 Mb region (352,091 markers) from 100k simulated UK-European reference samples into 1000, 100, and 10 target samples genotyped for 3,333 markers. Imputation analyses were run on a 2.6 GHz Intel Xeon E5-2630v2 computer with 128 GB of memory. CPU time is the sum of user and system time reported by the Unix time command. Relative CPU time is the ratio of the CPU time to the Beagle 5.0 CPU time for the same number of target samples. A dash (---) indicates the analysis could not be performed due to software limitations.

| Method | Target (n) | CPU seconds per target sample | Relative CPU time |
| --- | --- | --- | --- |
| **Beagle 5.0** | 1000 | 0.17 | 1.00 |
| **Beagle 4.1** | 1000 | 14.55 | 84.23 |
| **Minimac4** | 1000 | 2.10 | 12.14 |
| **Minimac3** | 1000 | 8.14 | 47.13 |
| **Impute4** | 1000 | --- | --- |
| **Beagle 5.0** | 100 | 0.29 | 1.00 |
| **Beagle 4.1** | 100 | 15.37 | 52.51 |
| **Minimac4** | 100 | 3.63 | 12.42 |
| **Minimac3** | 100 | 8.35 | 28.54 |
| **Impute4** | 100 | --- | --- |
| **Beagle 5.0** | 10 | 1.25 | 1.00 |
| **Beagle 4.1** | 10 | 24.77 | 19.76 |
| **Minimac4** | 10 | 17.41 | 13.89 |
| **Minimac3** | 10 | 10.45 | 8.33 |
| **Impute4** | 10 | --- | --- |

**Table S5:**
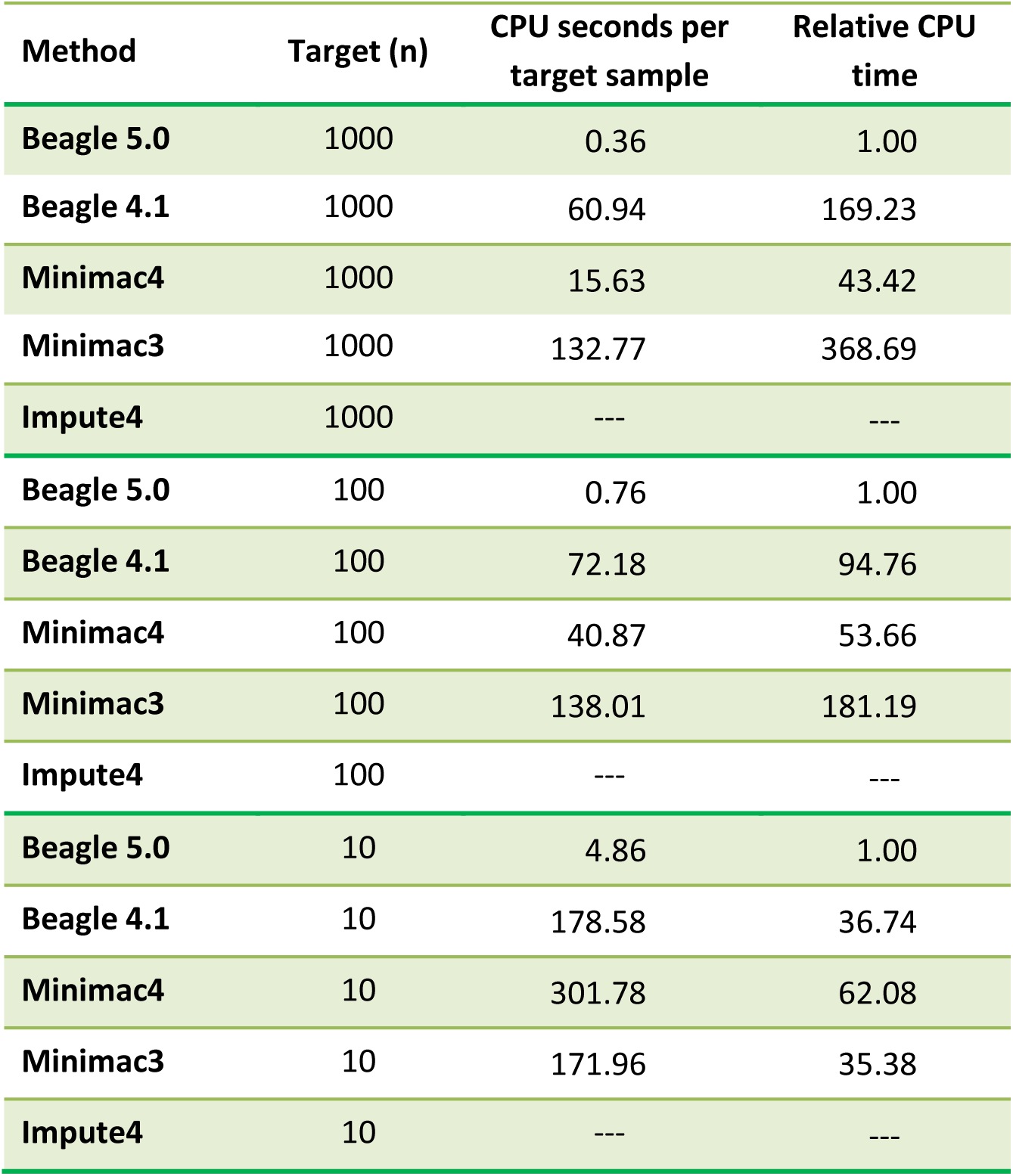
Single-threaded CPU time for imputation from 1M simulated UK-European reference samples. Single-threaded CPU time for Beagle 5.0, Beagle 4.1, Minimac4, Minimac3, and Impute4 to impute a 10 Mb region (1,210,904 markers) from 1M simulated UK-European reference samples into 1000, 100, and 10 target samples genotyped for 3,333 markers. Imputation analyses were run on a 2.6 GHz Intel Xeon E5-2630v2 computer with 128 GB of memory. CPU time is the sum of user and system time reported by the Unix time command. Relative CPU time is the ratio of the CPU time to the Beagle 5.0 CPU time for the same number of target samples. A dash (---) indicates the analysis could not be performed due to software limitations.

**Table S6:**
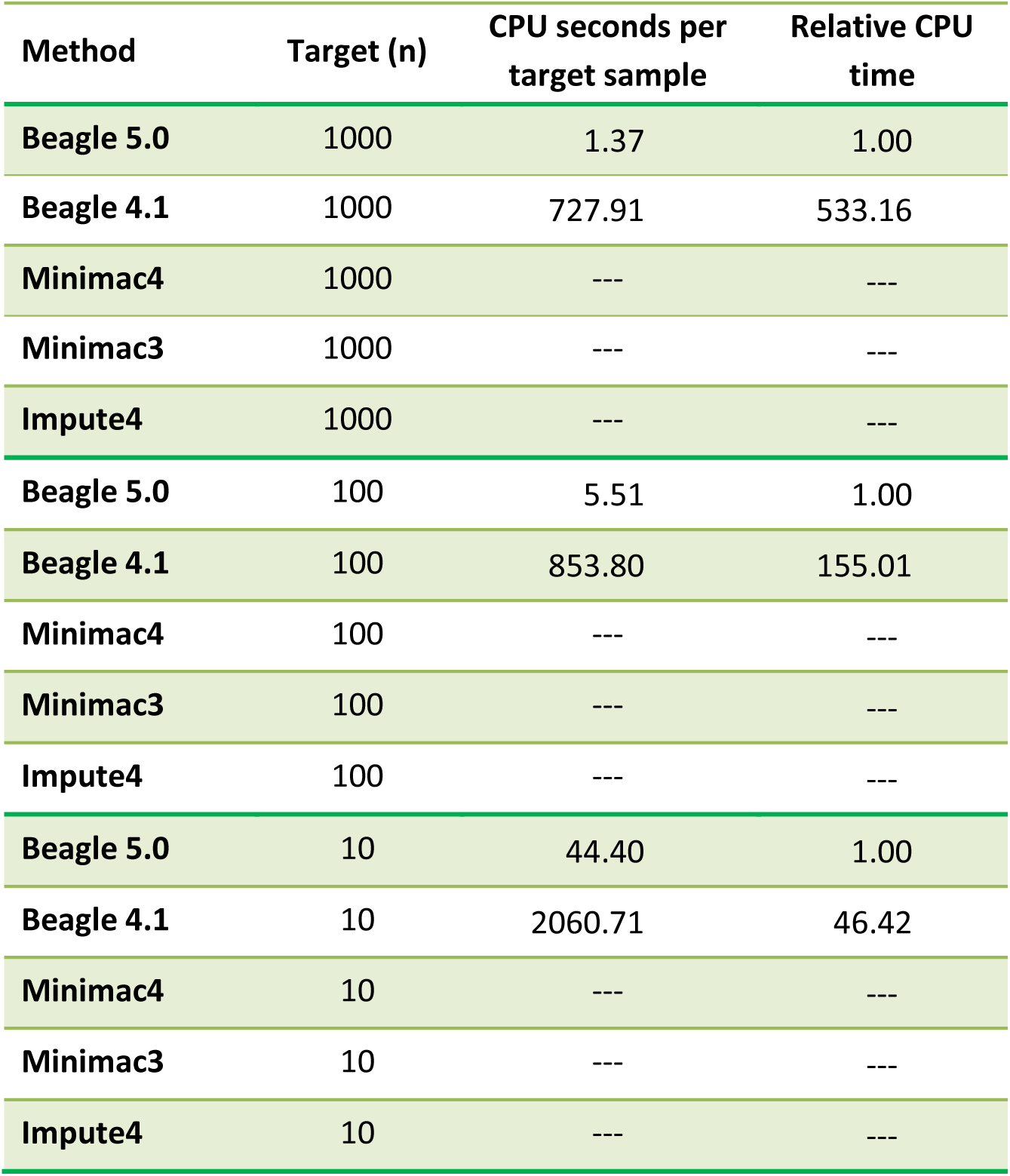
Single-threaded CPU time for imputation from 10M simulated UK-European reference samples. Single-threaded CPU time for Beagle 5.0, Beagle 4.1, Minimac4, Minimac3, and Impute4 when imputing a 10 Mb region (3,145,425 markers) from 1M simulated UK-European reference samples into 1000, 100, and 10 target samples genotyped for 3,333 markers. Imputation analyses were run on a 2.6 GHz Intel Xeon E5-2630v2 computer with 128 GB of memory. CPU time is the sum of user and system time reported by the Unix time command. Relative CPU time is the ratio of the CPU time to the Beagle 5.0 CPU time for the same number of target samples. A dash (---) indicates the analysis could not be performed due to memory constraints or software limitations.

**Table S7:**
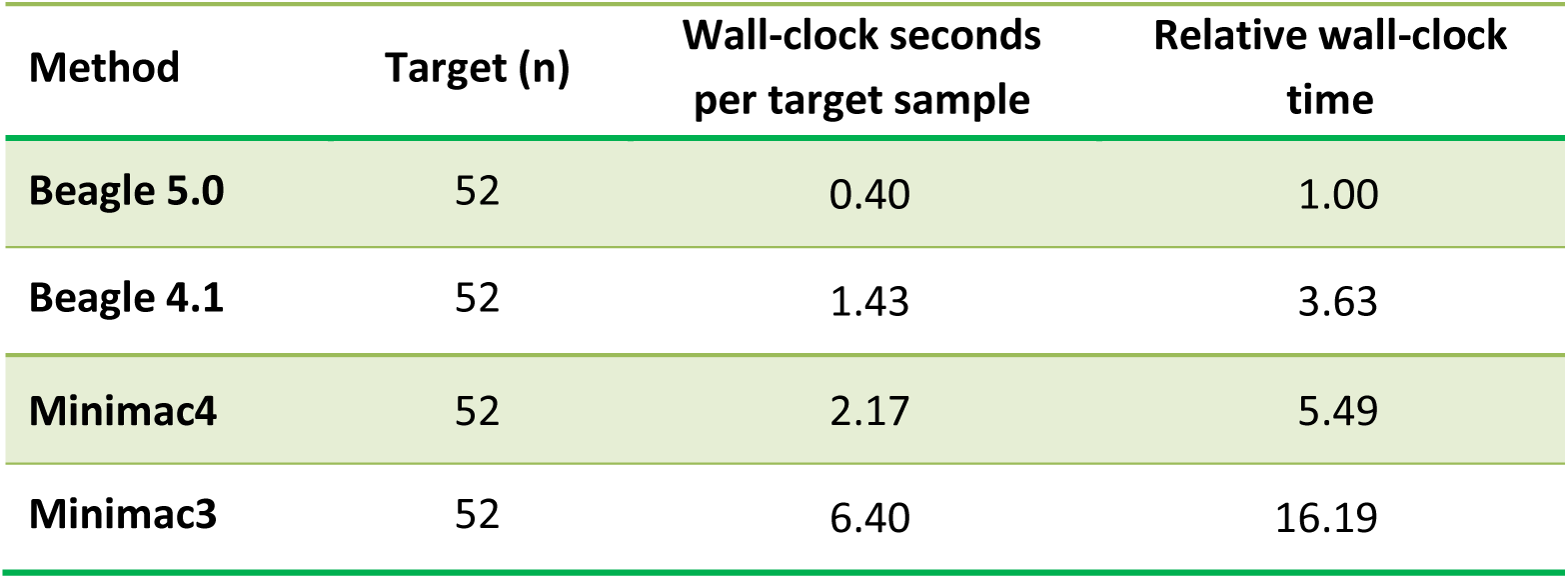
Multi-threaded wall-clock time for imputation from 2,452 reference samples from the 1000 Genomes Project. Multi-threaded wall-clock time for Beagle 5.0, Beagle 4.1, Minimac4, and Minimac3 when using 12 CPU cores to impute chromosome 20 (1,718,742 markers) from 2,452 reference samples from the 1000 Genomes Project into 52 target samples genotyped for 54,885 markers. Imputation analyses were run on a 12-core 2.6 GHz Intel Xeon E5-2630v2 computer with 128 GB of memory. Relative wall-clock time is the ratio of the wall-clock time to the Beagle 5.0 wall-clock time.

| Method | Target (n) | Wall-clock seconds<br>per target sample | Relative wall-clock<br>time |
| --- | --- | --- | --- |
| Beagle 5.0 | 52 | 0.40 | 1.00 |
| Beagle 4.1 | 52 | 1.43 | 3.63 |
| Minimac4 | 52 | 2.17 | 5.49 |
| Minimac3 | 52 | 6.40 | 16.19 |

**Table S8:** Multi-threaded wall-clock time for imputation from 26,165 Haplotype Reference Consortium reference samples. Multi-threaded wall-clock time for Beagle 5.0, Beagle 4.1, Minimac4, and Minimac3 when using 12 CPU cores to impute chromosome 20 (862,298 markers) from 26,165 Haplotype Reference Consortium reference samples into 1000, 100, and 10 target samples genotyped for 55,013 markers. Imputation analyses were run on a 12-core 2.6 GHz Intel Xeon E5-2630v2 computer with 128 GB of memory. Relative wall-clock time is the ratio of the wall-clock time to the Beagle 5.0 wall-clock time for the same number of target samples.

| Method | Target (n) | Wall-clock seconds per target sample | Relative wall-clock time |
| --- | --- | --- | --- |
| Beagle 5.0 | 1000 | 0.20 | 1.00 |
| Beagle 4.1 | 1000 | 4.75 | 23.24 |
| Minimac4 | 1000 | 2.29 | 11.19 |
| Minimac3 | 1000 | 8.67 | 42.45 |
| Beagle 5.0 | 100 | 0.48 | 1.00 |
| Beagle 4.1 | 100 | 6.76 | 14.15 |
| Minimac4 | 100 | 7.11 | 14.90 |
| Minimac3 | 100 | 9.38 | 19.65 |
| Beagle 5.0 | 10 | 3.28 | 1.00 |
| Beagle 4.1 | 10 | 24.48 | 7.48 |
| Minimac4 | 10 | 54.85 | 16.75 |
| Minimac3 | 10 | 13.22 | 4.04 |

**Table S9:** Multi-threaded wall-clock time for imputation from 10k simulated UK-European reference samples. Multi-threaded wall-clock time for Beagle 5.0, Beagle 4.1, Minimac4, and Minimac3 when using 12 CPU cores to impute a 10 Mb region (88,124 markers) from 10k simulated UK-European reference samples into 1000, 100, and 10 target samples genotyped for 3,333 markers. Imputation analyses were run on a 12-core 2.6 GHz Intel Xeon E5-2630v2 computer with 128 GB of memory. Relative wall-clock time is the ratio of the wall-clock time to the Beagle 5.0 wall-clock time for the same number of target samples.

| Method | Target (n) | Wall-clock seconds<br>per target sample | Relative wall-clock<br>time |
| --- | --- | --- | --- |
| Beagle 5.0 | 1000 | 0.01 | 1.00 |
| Beagle 4.1 | 1000 | 0.14 | 9.70 |
| Minimac4 | 1000 | 0.07 | 5.15 |
| Minimac3 | 1000 | 0.25 | 17.39 |
| Beagle 5.0 | 100 | 0.05 | 1.00 |
| Beagle 4.1 | 100 | 0.21 | 4.62 |
| Minimac4 | 100 | 0.18 | 3.91 |
| Minimac3 | 100 | 0.32 | 6.83 |
| Beagle 5.0 | 10 | 0.32 | 1.00 |
| Beagle 4.1 | 10 | 0.82 | 2.58 |
| Minimac4 | 10 | 1.21 | 3.79 |
| Minimac3 | 10 | 0.86 | 2.71 |

**Table S10:** Multi-threaded wall-clock time for imputation from 100k simulated UK-European reference samples. Multi-threaded wall-clock time for Beagle 5.0, Beagle 4.1, Minimac4, and Minimac3 when using 12 CPU cores to impute a 10 Mb region (352,091 markers) from 100k simulated UK-European reference samples into 1000, 100, and 10 target samples genotyped for 3,333 markers. Imputation analyses were run on a 12-core 2.6 GHz Intel Xeon E5-2630v2 computer with 128 GB of memory. Relative wall-clock time is the ratio of the wall-clock time to the Beagle 5.0 wall-clock time for the same number of target samples.

| Method | Target (n) | Wall-clock seconds per target sample | Relative wall-clock time |
| --- | --- | --- | --- |
| Beagle 5.0 | 1000 | 0.02 | 1.00 |
| Beagle 4.1 | 1000 | 1.34 | 66.07 |
| Minimac4 | 1000 | 0.46 | 22.74 |
| Minimac3 | 1000 | 4.31 | 212.06 |
| Beagle 5.0 | 100 | 0.06 | 1.00 |
| Beagle 4.1 | 100 | 1.82 | 28.84 |
| Minimac4 | 100 | 1.86 | 29.54 |
| Minimac3 | 100 | 4.53 | 72.00 |
| Beagle 5.0 | 10 | 0.46 | 1.00 |
| Beagle 4.1 | 10 | 5.87 | 12.66 |
| Minimac4 | 10 | 15.54 | 33.52 |
| Minimac3 | 10 | 6.81 | 14.70 |

**Table S11:**
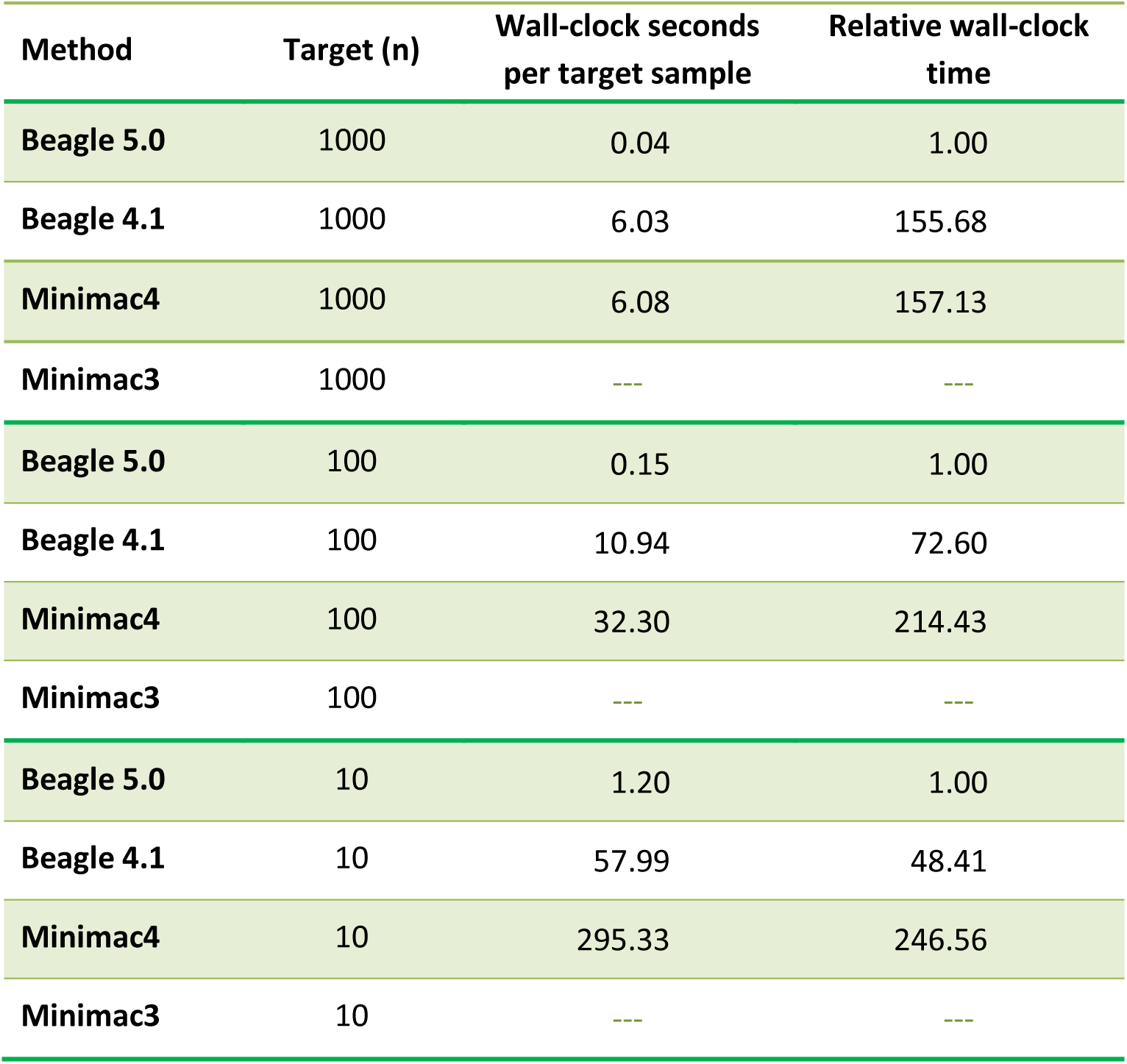
Multi-threaded wall-clock time for imputation from 1M simulated UK-European reference samples. Multi-threaded wall-clock time for Beagle 5.0, Beagle 4.1, Minimac4, and Minimac3 when using 12 CPU cores to impute a 10 Mb region (1,210,904 markers) from 1M simulated UK-European reference samples into 1000, 100, and 10 target samples genotyped for 3,333 markers. Imputation analyses were run on a 12-core 2.6 GHz Intel Xeon E5-2630v2 computer with 128 GB of memory. Relative wall-clock time is the ratio of the wall-clock time to the Beagle 5.0 wall-clock time for the same number of target samples. A dash (---) indicates the analysis could not be performed due to memory constraints.

| Method | Target (n) | Wall-clock seconds per target sample | Relative wall-clock time |
| --- | --- | --- | --- |
| Beagle 5.0 | 1000 | 0.04 | 1.00 |
| Beagle 4.1 | 1000 | 6.03 | 155.68 |
| Minimac4 | 1000 | 6.08 | 157.13 |
| Minimac3 | 1000 | --- | --- |
| Beagle 5.0 | 100 | 0.15 | 1.00 |
| Beagle 4.1 | 100 | 10.94 | 72.60 |
| Minimac4 | 100 | 32.30 | 214.43 |
| Minimac3 | 100 | --- | --- |
| Beagle 5.0 | 10 | 1.20 | 1.00 |
| Beagle 4.1 | 10 | 57.99 | 48.41 |
| Minimac4 | 10 | 295.33 | 246.56 |
| Minimac3 | 10 | --- | --- |

**Table S12:** Multi-threaded wall-clock time for imputation from 1GM simulated UK-European reference samples. Multi-threaded wall-clock time for Beagle 5.0, Beagle 4.1, Minimac4, and Minimac3 when using 12 CPU cores to impute a 10 Mb region (3,145,425 markers) from 10M simulated UK-European reference samples into 1000, 100, and 10 target samples genotyped for 3,333 markers. Imputation analyses were run on a 12-core 2.6 GHz Intel Xeon E5-2630v2 computer with 128 GB of memory. Relative wall-clock time is the ratio of the wall-clock time to the Beagle 5.0 wall-clock time for the same number of target samples. A dash (---) indicates the analysis could not be performed due to memory constraints.

| Method | Target (n) | Wall-clock seconds<br>per target sample | Relative wall-clock<br>time |
| --- | --- | --- | --- |
| Beagle 5.0 | 1000 | 0.16 | 1.00 |
| Beagle 4.1 | 1000 | 72.83 | 457.99 |
| Minimac4 | 1000 | --- | --- |
| Minimac3 | 1000 | --- | --- |
| Beagle 5.0 | 100 | 0.83 | 1.00 |
| Beagle 4.1 | 100 | 139.65 | 167.54 |
| Minimac4 | 100 | --- | --- |
| Minimac3 | 100 | --- | --- |
| Beagle 5.0 | 10 | 7.12 | 1.00 |
| Beagle 4.1 | 10 | 662.04 | 93.00 |
| Minimac4 | 10 | --- | --- |
| Minimac3 | 10 | --- | --- |

**Figure S13:**
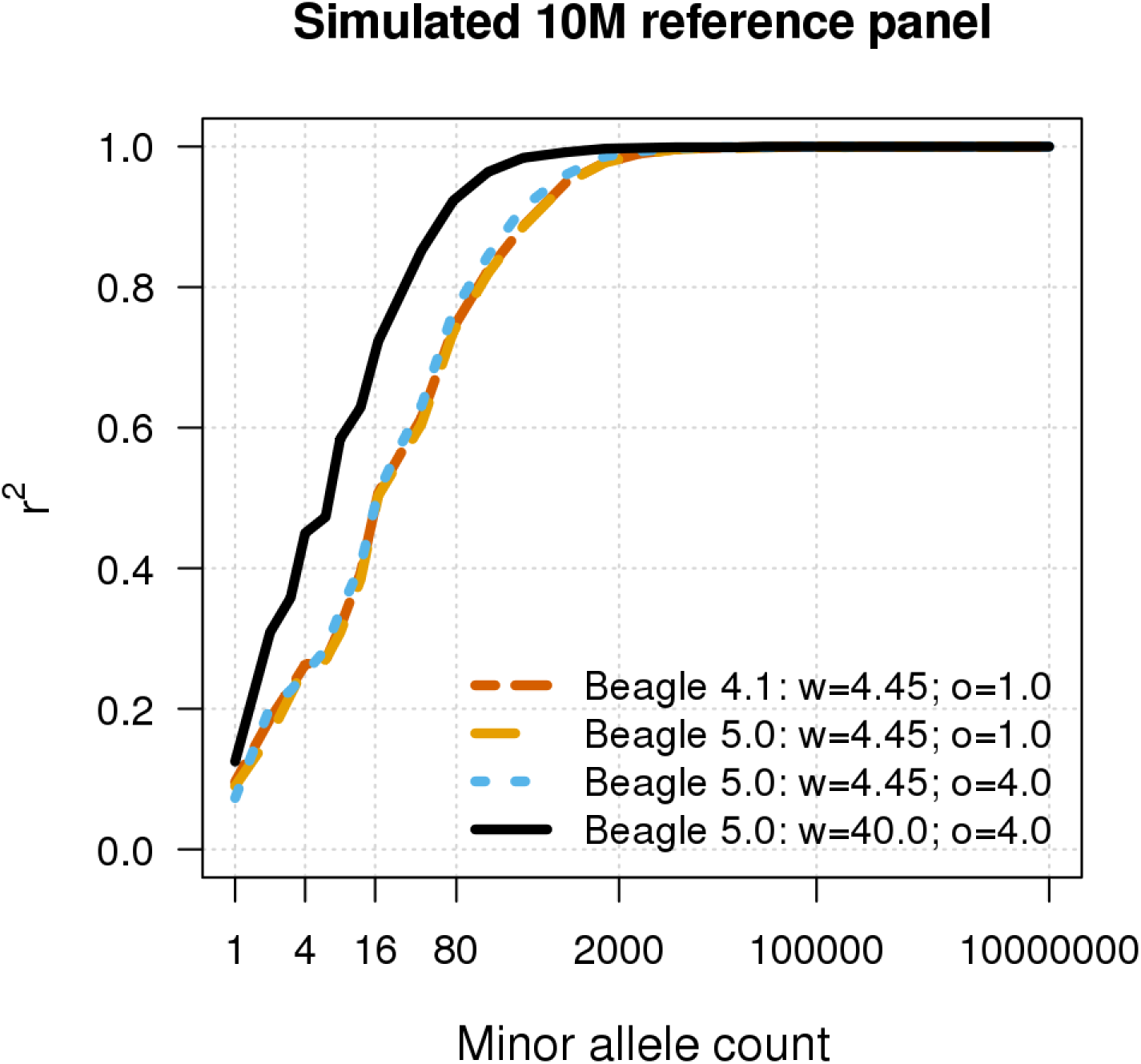
Effect of window length and overlap on imputation accuracy. Genotype imputation accuracy for Beagle 5.0 and Beagle 4.1 when imputing genotypes from reference panels with 10M simulated UK-European reference samples. Imputed alleles are binned according to their minor allele count in each reference panel. The squared correlation (*r^2^*) between the true number of alleles on a haplotype (0 or 1) and the imputed posterior allele probability is reported for each minor allele count bin. The horizontal axis in each panel is on a log scale. Beagle 4.1 was run with a 4.45 cM window and 1.0 cM overlap (w=4.45; o=1.0). Beagle 5.0 was run with three different combinations of window length and overlap: 1) 4.45 cM window and 1.0 cM overlap (w=4.45; o=1.0), 2) 4.45 cM window and 4.0 cM overlap (w=4.45; o=4.0), and 3) 10.0 cM window and 4.0 cM overlap (w=10.0; o=1.0).

## Text S14

### Msprime script for simulating UK-European samples

~~~
# Msprime simulation script for generating a user-specified length of
# chromosome with constant recombination rate for UK individuals with
# European ancestry.
# Population history for eur, yri, asn is based on the Gravel et al.
# 2011 analysis (see https://doi.org/10.1073/pnas.1019276108).
# Recent growth is based on our IBDNe analysis of the UK10K data
# (see https://doi.org/10.1016/j.ajhg.2015.07.012).
# Growth rates change at 150 generations ago (due to introduction
# of agriculture) and 15 generations ago (start of increased growth
# seen in IBDNe analysis).
# Fitted effective population size is 3.96e6 now, 5.64e5 at 15
# generations ago, 9.80e3 at 150 generations ago. Hence growth rate
#.00292 until 150 generations ago, then .030 until 15 generations
# ago, then .13 until present
# Msprime generates positions of mutations as floating numbers, but
# converts to integer positions when writing the vcf file, and it
# outputs only diallelic variants. Our simulation script rescales
# distance so that distinct mutations occur at distinct positions.
# After simulation, the output positions should be rescaled by
# dividing by the scale factor, and mutations mapping to the same
# rescaled position can be converted into multi-allelic markers.
# usage: python2.7 uk_scale.py seed nhaps nbp scale > vcf.file
# for the results in the paper, we used seed=1; nhaps=20002000;
# nbp=10000000; scale=100.
import msprime, sys
from math import log
from math import exp
seed = int(sys.argv[1])
nhaps = int(sys.argv[2])
scale = int(sys.argv[4])
nbp = int(sys.argv[3])*scale
mu=1.25e-8/scale # mutation rate per bp
rho=1e-8/scale # recombination rate per bp
genlen=25 # years per generation
N0=7310 # initial population size, and reference effective size
Thum=5920 # =148000/genlen; generations back to initial pop growth (advent of modern humans)
Naf=14474 # size of african population from t0 onwards
Tooa=2040 #51000/genlen # number of generations back to Out of Africa
Nb=1861                            # size of out of Africa population
mafb=1.5e-4           # migration rate between Africa and Out-of-Africa
Teu=920 #23000/genlen # number of generations back to Asia-Europe split
Neu=1032; Nas=554 # bottleneck population sizes after the split
mafeu=2.5e-5; mafas=7.8e-6; meuas=3.11e-5 # migration rates between Africa, Europe and Asia
ras=0.0048 # growth rates per generation in Asia
reu=0.00292
Tex=150    # generations back to accelerated population growth
rex=0.03    # accelerated growth rate
Tmod=15
rmod=0.13 # growth rate in most recent generations
# pop0 is Africa, pop1 is eur (uk), pop2 is east asn
# will generate nhaps from eur pop
samplesize = nhaps
othersize = 0
pop_config = [
      msprime.PopulationConfiguration(sample_size=samplesize,initial_size= Neu*exp(reu*(Teu-Tex))*
exp(rex*(Tex-Tmod))* exp(rmod*Tmod), growth_rate=rmod),
      msprime.PopulationConfiguration(sample_size=othersize,initial_size= Nas*exp((Teu-Tex)*ras)*
exp(rex*Tex),growth_rate=rex)
]
mig_mat = [[0,mafeu,mafas],[mafeu,0,meuas],[mafas,meuas,0]]
# recent change in growth rate
recent_event = [
      msprime.PopulationParametersChange(time=Tmod,growth_rate=rex,population_id=1)
]
# populations stop having accelerated growth (advent of agriculture)
ag_event = [
      msprime.PopulationParametersChange(time=Tex,growth_rate=ras,population_id=2),
      msprime.PopulationParametersChange(time=Tex,growth_rate=reu,population_id=1),
      msprime.PopulationParametersChange(time=Tex,growth_rate=0.0,population_id=0)
]
# Asia and Europe merge, migration changes, population size changes, growth stops
eu_event = [
      msprime.MigrationRateChange(time=Teu,rate=0.0),
      msprime.PopulationParametersChange(time=Teu,growth_rate=0.0, population_id=2),
      msprime.MassMigration(time=Teu+0.0001,source=2,destination=1,proportion=1.0),
      msprime.PopulationParametersChange(time=Teu+0.0001,initial_size=Nb,growth_rate=0.0, population_id=1),
      msprime.MigrationRateChange(time=Teu+0.0001,rate=mafb,matrix_index=(0,1)),
      msprime.MigrationRateChange(time=Teu+0.0001,rate=mafb,matrix_index=(1,0))
]
# Out of Africa event (looking back, Africa and Europe merge)
ooa_event = [
      msprime.MigrationRateChange(time=Tooa,rate=0.0),
      msprime.MassMigration(time=Tooa+0.0001,source=1,destination=0,proportion=1.0)
]
# initial population size
init_event = [
      msprime.PopulationParametersChange(time=Thum,initial_size=N0,population_id=0)
]
# cat all the events together
events = recent_event + ag_event + eu_event + ooa_event + init_event
# run the simulation
treeseq = msprime.simulate(population_configurations=pop_config, migration_matrix=mig_mat, demographic_events=events, length=nbp, recombination_rate=rho, mutation_rate=mu, random_seed=seed)
# print results
with sys.stdout as vcffile:
              treeseq.write_vcf(vcffile,2) # 2 is for diploid
~~~

